# Divergent neural pathways emanating from the lateral parabrachial nucleus mediate distinct components of the pain response

**DOI:** 10.1101/602466

**Authors:** Michael C. Chiang, Eileen K. Nguyen, Andrew E. Papale, Sarah E. Ross

## Abstract

The lateral parabrachial nucleus (lPBN) is a major target of spinal projection neurons conveying nociceptive input into supraspinal structures. However, the functional role of distinct lPBN efferents for diverse nocifensive responses have remained largely uncharacterized. Here, we show that two populations of efferent neurons from different regions of the lPBN collateralize to distinct targets. Activation of efferent projections to the ventromedial hypothalamus (VMH) or lateral periaqueductal gray (lPAG) drive escape behaviors, whereas the activation of lPBN efferents to the bed nucleus stria terminalis (BNST) or central amygdala (CEA) generates an aversive memory. Finally, we provide evidence that dynorphin expressing neurons span cytoarchitecturally distinct domains of the lPBN to coordinate these distinct aspects of the nocifensive response.

**HIGHLIGHTS:** Spatially segregated neurons in the lPBN collateralize to distinct targets.

Distinct output pathways give rise to separate aspects of the pain response.

Dynorphin neurons within the lPBN convey noxious information across subdivisions.

**eTOC BLURB:** Chiang et al. reveal that neurons in spatially segregated regions of the lateral parabrachial nucleus collateralize to distinct targets, and that activation of distinct efferents gives rise to separate components of the nocifensive response.

## INTRODUCTION

Noxious stimuli elicit a repertoire of innately encoded nocifensive behaviors comprising locomotor actions and affective responses to prevent further potential damage. These stereotyped behavioral sequences shift toward more complex motor actions as the severity of potential damage increases. For example, low intensity noxious stimuli drive withdrawal reflexes, whereas high intensity noxious stimuli recruit escape behaviors and aversive learning. Together, these distinct behavioral responses form a nocifensive response (Browne et al., 2017; Espejo and Mir, 1993; Fan et al., 1995; Le Bars et al., 2001). Despite the importance of the appropriate response for survival, the neural underpinnings of the distinct components that make up the nocifensive response remains to be fully explored.

It is well established that the lateral parabrachial nucleus (lPBN) is a primary target for nociceptive information arising from the spinal cord (Al-Khater and Todd, 2009; Todd et al., 2000). Indeed, the majority of lPBN neurons respond to noxious stimuli ((Bester et al., 1997; Hermanson and Blomqvist, 1996, 1997; Jansen and Giesler, 2015; Menendez et al., 1996). However, the lPBN is not specific to nociception, as it is activated by a wide array of threatening stimuli, such as food neophobia and hypercapnia (Chamberlin and Saper, 1994; Kaur et al., 2013; Kaur et al., 2017; Palmiter, 2018; Saper, 2016). Recently, the contribution of a specific population of lPBN neurons, those that express calcitonin gene-related peptide (CGRP) neurons, were found to play an important role in fear learning via projections to the central amygdala (Campos et al., 2018; Han et al., 2015). However, these CGRP neurons represent only a portion of the projections from the lPBN. Given that so many lPBN neurons respond to noxious stimulation, we sought to gain a clearer understanding of its efferents and how their activity might contribute to the response to noxious stimuli.

In this study, we investigated the varying contributions of distinct lPBN efferents to the the bed nucleus stria terminalis (BNST), central amygdala (CEA), ventromedial hypothalamus (VMH), and lateral periaqueductal gray (lPAG). We found that subsets of neurons in spatially segregated regions within the lPBN collateralize to distinct targets. Optogenetic manipulation of these specific outputs recapitulates specific components of a nocifensive response. Furthermore, we characterize a previously unspecified local lPBN circuit involving dynorphin neurons that are activated by noxious stimuli and may convey this information across lPBN subdivisions to mediate aversion.

## RESULTS

Nociceptive information is conveyed from the spinal cord to multiple regions of the brain in parallel, including brainstem, midbrain and forebrain structures (Todd, 2010). Although the lPBN is a major target of the anterolateral tract in murine species (Cameron et al., 2015; Todd, 2010; Todd et al., 2000), its relative contribution to pain behaviors has only recently been explored (Alhadeff et al., 2018; Barik et al., 2018; Huang et al., 2018; Rodriguez et al., 2017). To further address this issue, we tested whether transiently inhibiting the lPBN would affect the behavioral response to noxious stimuli. For these experiments, adenoassociated virus (AAV) encoding a Cre-dependent channelrhodopsin2 (ChR2) or enhanced yellow fluorescent protein (eYFP) was targeted into inhibitory neurons in lPBN using the *Gad2^cre^* allele to enable light-activated inhibition (Figures 1A and 1B). In the absence of light, both ChR2 and eYFP mice showed capsaicin-induced mechanical hypersensitivity. However, this hypersensitivity was significantly reduced when the lPBN was photostimulated in ChR2 mice compared to eYFP controls (Figure 1C). These data suggest that activity within the lPBN is required for mechanical allodynia.

**Figure 1.**
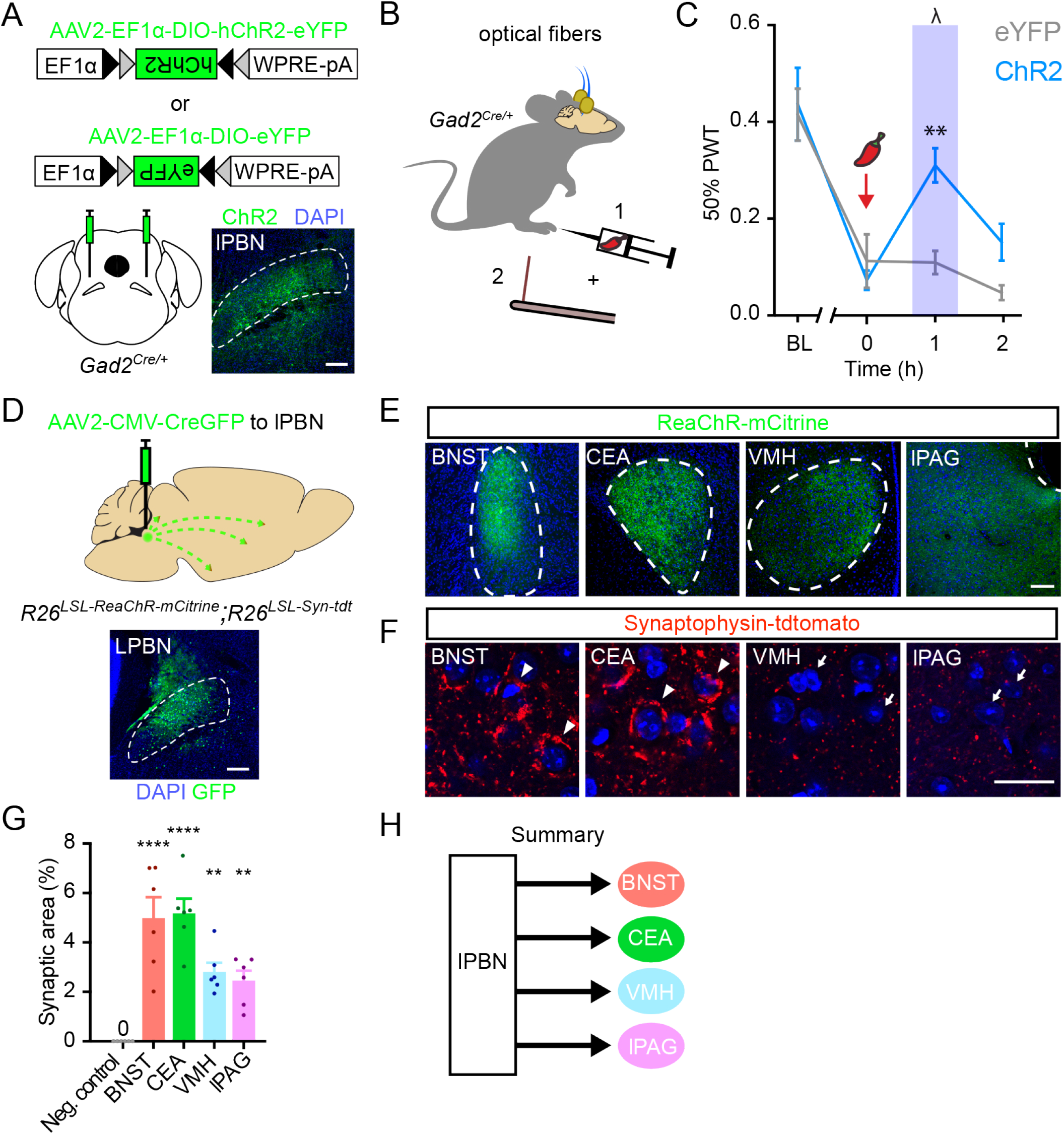
lPBN is required for mechanical hypersensitivity and has four major efferent targets. (A) Experimental strategy to drive inhibition in the lPBN. AAVs encoding Cre-dependent ChR2 or eYFP were bilaterally injected into the lPBN of *Gad2^cre^* mice. Representative image depicts expression of ChR2 within the lPBN (outline). Scale bar = 100 μm. (B) Mechanical hypersensitivity was (1) induced through intraplantar injection of capsaicin (10 μl, 0.03%) and (2) tested using von Frey filaments. (C) Paw withdrawal threshold (PWT) was significantly reduced during optogenetic stimulation (blue bar) in ChR2-expressing mice compared to eYFP-expressing controls. Data are mean ± SEM (n = 10 – 11 mice per group) ** indicates significantly different (two-way RM ANOVA followed by Holm-Sidak post-hoc test: p = 0.008). (D) Experimental design and representative image of strategy to visualize lPBN neurons, their projections, and their presynaptic terminals. An AAV encoding a Cre-GFP fusion protein was targeted to the lPBN into mice harboring two Cre-dependent alleles, *R26^LSL-ReaChR-mCitrine^* and *R26^LSL-Syn-tdt^*. Scale bar = 100 μm. (E) Projections of the lPBN efferents to four different brain regions, as visualized with ReaChR-mCitrine: BSNT, CEA, VMH and lPAG. Scale bar = 100 μm. Images are representative of results from 6 mice. (F) Synaptic terminals of lPBN efferents at four indicated targets, as visualized with Synaptophysin-tdtomato. Scale bar = 25 μm. Arrowheads and arrows denote perisomatic and diffuse input, respectively. (G) Quantification of synaptic input. The relative number of synapses from lPBN was estimated by quantifying the area of synaptophysin-tdtomato expression within the indicated target. An arbitrary brain region with no synaptophysin-tdtomato expression was used as negative control. Data are mean ± SEM and dots represent data points from individual animals (n = 6 mice). Asterisks indicate significantly different than negative control region (one-way RM ANOVA followed by Holm-Sidak post-hoc test; ** p < 0.01, **** p < 0.001). (H) Summary diagram depicting four major efferent targets of lPBN.

Given the necessity of the lPBN for this pain behavior, we next explored its efferent targets. Towards this end, an AAV encoding a Cre-GFP fusion protein was stereotaxically delivered into the lPBN of mice harboring two Cre-dependent alleles: ReaChR-mCitrine, for the purpose of visualizing axonal projections, and synaptophysin-tdTomato, for the purpose of visualizing presynaptic terminals (Figure 1D). We observed lPBN efferent projections to numerous regions of the brain (Figure S1), consistent with previous studies (Bernard et al., 1996; Bernard et al., 1994; Gauriau and Bernard, 2002; Saper and Loewy, 1980). However, four targets in particular stood out due to the robust projections from mCitrine-labeled axons and dense puncta from tdtomato-labeled synaptic terminals: the BNST, the CEA, the VMH, and the lPAG (Figures 1E and 1F). Quantification revealed that all four of these regions received significant synaptic input from the lPBN (Figure 1G), though the apparent perisomatic input to the BNST and CEA (arrows) was qualitatively different from the diffuse input observed within the VMH and lPAG (arrowheads; Figure 1F). Together, these data indicate that the BNST, CEA, VMH, and lPAG are four principle efferent targets of the lPBN (Figure 1H).

Next, we sought to investigate the cellular basis of these efferent projections in more detail. In particular, we considered whether there might be parallel pathways originating from distinct cell types within the lPBN (Figure 2A), which would be consistent with previous work suggesting that distinct subdivisions of the LBPN have distinct projection patterns (Fulwiler and Saper, 1984; Saper and Loewy, 1980). Alternatively, given that at least some lPBN efferents are known to collateralize (Tokita et al., 2010), we also considered the possibility of a single major output from the lPBN with multiple targets (Figure 2B). To distinguish between these possibilities, we characterized the projections from the lPBN using cholera toxin B subunit (CTB) as a retrograde tracing tool (Figure 2C). Intriguingly, we found that stereotaxic injection of CTB into distinct lPBN targets labeled neuronal cell bodies in different sub-regions of the lPBN: retrograde tracing from BNST or CEA resulted in labeled neurons within the external lateral division (elPBN), whereas retrograde tracing from VMH or lPAG labeled neurons within the dorsal division (dPBN) (Figure 2D). These findings suggested the existence of at least two populations of efferent neurons with distinct targets.

**Figure 2.**
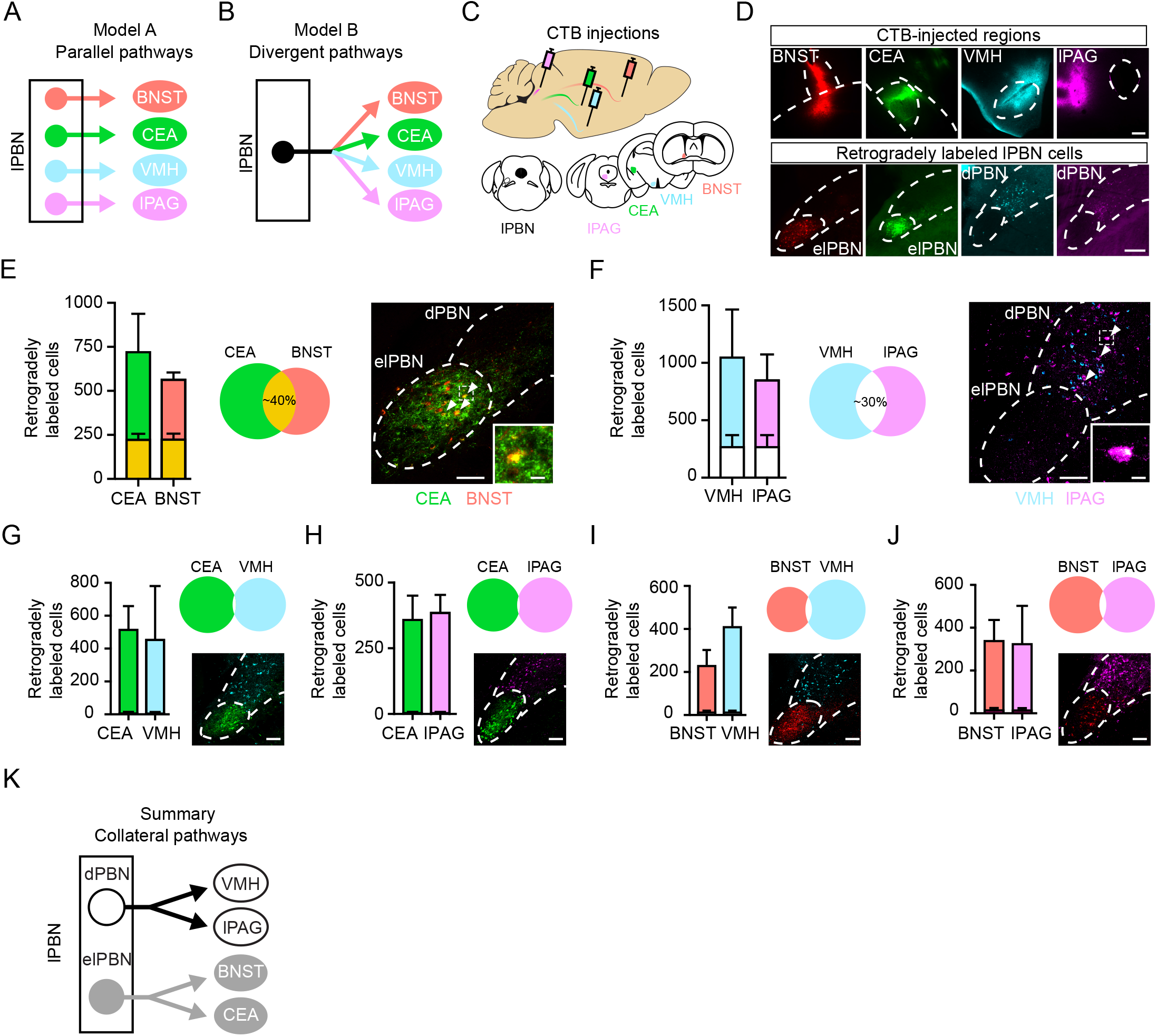
Distinct subpopulations of lPBN collateralize to different forebrain regions. (A-B) Models illustrating lPBN efferents as parallel (A) or divergent pathways (B) (C) Experimental strategy to retrogradely label lPBN efferents with fluorophore-congugated CTB. (D) CTB injections into efferent targets (top) and retrogradely labeled cells (bottom) in elPBN (BSNT and CEA) and dPBN (VMH and lPAG). Scale bars = 100 μm. (E) Dual injection of CTB into CEA (green) and BNST (red) resulted in colocalized signal in approximately 40% of retrogradely labeled cells (yellow) across entire lPBN. Data are mean ± SEM (n = 4 mice). Arrows highlight co-labeled cells. Scale bar = 50 μm. Magnification shown in inset. Scale bar = 10 μm. (F) Dual injections of CTB into VMH (blue) and lPAG (purple) resulted in colocalized signal in 30% of retrogradely labeled cells (white) across entire lPBN. Data are mean ± SEM (n = 4 mice). Arrows highlight co-labeled cells. Scale bar = 50 μm. Magnification shown in inset. Scale bar = 10 μm. (G-J) Very few dual-labeled neurons were observed following dual CTB injections into: CEA and VMH (G); CEA and lPAG (H); BNST and VMH (I); or BNST and lPAG (J). Data are mean ± SEM (n = 3 - 4 mice). Scale bar = 50 μm. (K) Summary diagram illustrating two collateral pathways emerging from lPBN.

To further explore this idea, we performed dual retrograde labeling experiments, placing distinct CTB conjugates into different target regions through stereotaxic injections. Following dual targeting of CEA and BNST, we found that ∼40% of labeled neurons in the lPBN were double-labeled with both CTB-conjugated fluorophores (Figure 2E). Analogously, following dual injection into the VMH and lPAG, ∼30% of CTB-containing neurons in the lPBN were double-labeled (Figure 2F). In contrast, there was almost no double labeling of lPBN neurons upon dual injections into any of the other four pair-wise combinations (Figures 2G, 2H, 2I and 2J). Together, these data define two major efferent pathways from the lPBN: one originating from the dPBN that collateralizes to the VMH and lPAG, and a second arising from the elPBN that collateralizes to the BNST and CEA (Figure 2K).

In light of these findings, we next determined whether distinct outputs from the lPBN mediate different components of the nocifensive response. To address this question, we targeted the lPBN with AAVs encoding either ChR2 or eYFP and implanted optical fibers above distinct efferent targets, thereby enabling pathway-selective stimulation (Figure 3A). For each mouse, behavioral experiments and post-hoc analysis of tissue for infection specificity in the lPBN and optical placement over the efferent target were performed in a blinded manner. In addition, pathway specificity was confirmed by analyzing the induction of Fos, a marker of neuronal activity, in response to optogenetic stimulation (Figure S2).

**Figure 3.**
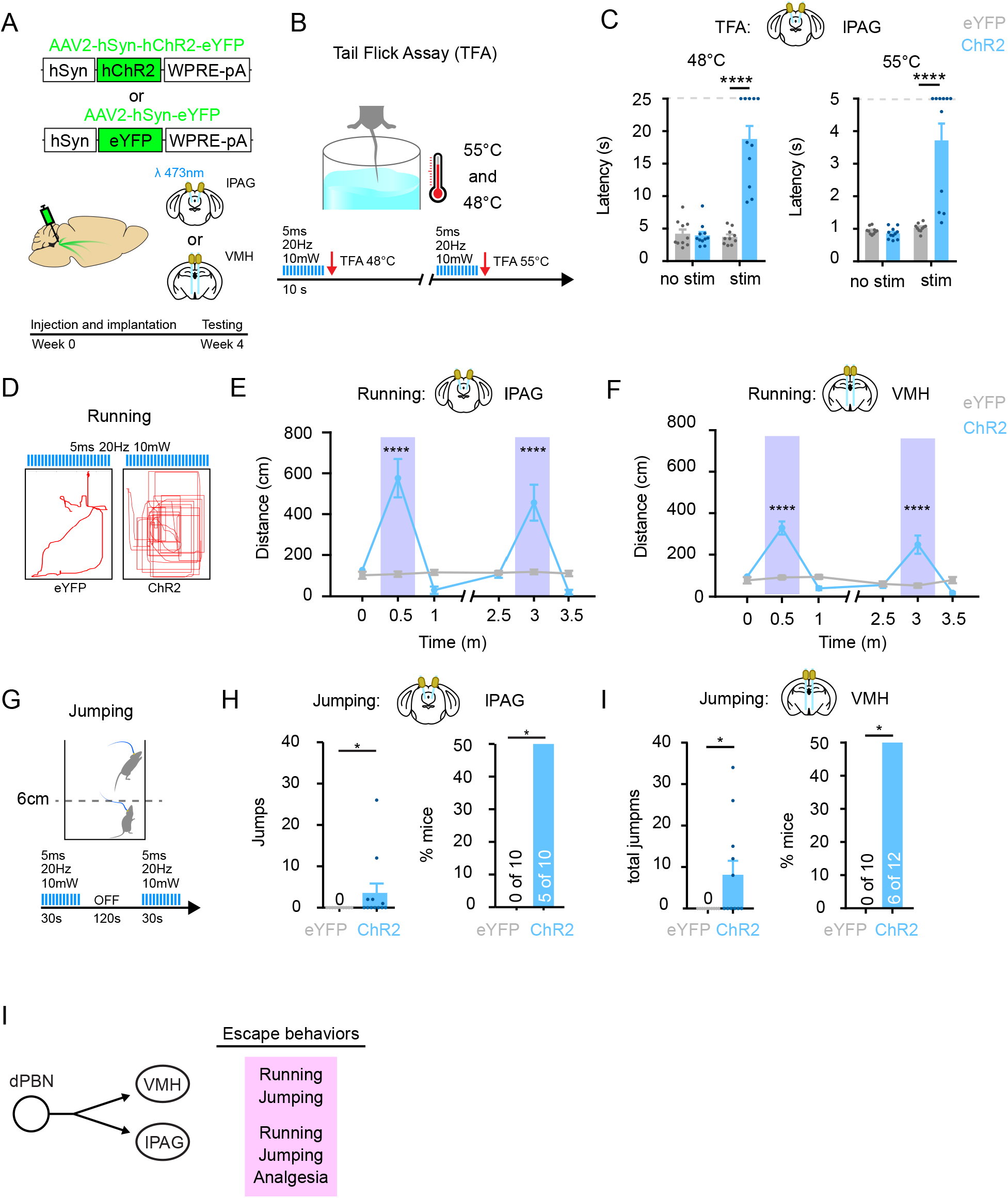
Efferent dPBN projections to VMH and lPAG elicit escape-like behaviors. (A) Experimental strategy to selectively activate distinct lPBN projections. AAVs encoding either ChR2 or eYFP were injected into the lPBN and optical implants were placed above one of four efferent targets: lPAG, VMH, CEA or BNST. (B) Experimental protocol for tail flick assay (TFA). Mice were photostimulated for 10 s immediately prior to TFA at either 48 °C and 55 °C. (C) Photostimulation of dPBN terminals in lPAG significantly increases latency to tail flick at 48 °C and 55 °C. Data are mean ± SEM and dots represent data points from individual animals (n = 9 – 11 mice per group). **** indicates significantly different (Two-way RM ANOVA followed by Holm-Sidak post-hoc test; lPAG, p < 0.0001). Dotted lines indicate cut-off latencies that were imposed to prevent tissue damage. (D) Experimental protocol for running assay. Stimulation paradigm and example traces of locomotion following stimulation of lPAG terminals from lPBN efferents in eYFP and ChR2 mice. (E) Photostimulation of dPBN terminals in lPAG significantly increases locomotion. Data are mean ± SEM (n = 9 – 11 mice per group). **** indicates significantly different (Two-way ANOVA followed by Holm-Sidak post-hoc test, p < 0.0001) (F) Photostimulation of dPBN terminals in VMH significantly increases locomotion. Data are mean ± SEM (n = 10 – 12 mice per group). **** indicates significantly different (Two-way ANOVA followed by Holm-Sidak post-hoc test, p < 0.0001). (G) Experimental protocol for jumping assay. A minimum of 6 cm vertical movement of the body was considered a jump. (H) Photostimulation of dPBN terminals in lPAG elicits significant jumping. Data are mean ± SEM and dots represent data points from individual animals (n = 9 – 11 mice per group). Left: * indicates significant number of jumps (Mann-Whitney; p = 0.033). Right: * indicates significant proportion of mice (Fisher’s exact test: p = 0.033). (I) Photostimulation of dPBN terminals in VMH elicits significant jumping. Data are mean ± SEM and dots represent data points from individual animals (n = 9 – 11 mice per group). Left: * indicates significant number of jumps (Mann-Whitney; p = 0.015). Right: * indicates significant proportion of mice (Fisher’s exact test: p = 0.015). (J) Summary diagram depicting behavioral responses observed upon stimulation of dPBN efferents to VMH and lPAG.

Several lines of evidence suggest that nociceptive threshold is determined, at least in part, by decending modulation from brain structures such as the PAG that are activated by ascending nociceptive circuitry (Basbaum and Fields, 1978). To explore whether any of the efferent projections from the lPBN are sufficient to activate descending inhibition, we assessed whether optogenetic stimulation affected the latency to withdraw in the tail flick assay, which measures a spinal reflex to noxious heat (Figure 3B). At baseline, ChR2-expressing mice exhibited similar tail flick latencies compared to eYFP controls. However, immediately following optogenetic activation of dPBN projections to lPAG, ChR2-expressing mice showed a significant increase in tail flick latency (Figure 3C). Indeed, over half of these mice reached cut-off, which was imposed to prevent tissue damage. In contrast, photostimulation of projections to other efferent targets had either no significant effect (VMH or CEA) or only a small effect (BNST) (Figures S3A, S3B and S3C). Thus, activation of the efferent pathway from the lPBN to the lPAG is sufficient to elicit robust analgesia through descending inhibition.

Over the course of these studies, we noted that activation of some efferent pathways elicited motor behaviors. To examine this phenomenon in more detail, we quantified the lateral (Figure 2D; running) and vertical (Figure 2G; jumping) movements that were observed upon optogenetic stimulation. Activation of the efferent projection from the dPBN to the lPAG resulted in explosive running behavior that was time-locked to the light stimulus (Figure 3E). Likewise, stimulation of the projection to the VMH elicited dramatic increases in locomotion that began each time the light was turned on and ceased as soon as the light was turned off (Figure 3F). In contrast, photostimulation of efferent projections to the CEA caused no significant lateral movement (Figure S3D), and that to the BNST showed significant lateral movement to the first stimulation only (Figure S2E). Thus, efferent projections from the dPBN were distinctive in their ability to elicit switch-like locomotor behavior in response to repeated stimulation.

Analogous results were found in the jumping assay, where significant effects were observed upon activation of efferents originating from the dPBN, but not the elPBN. Upon activation of projections to either the lPAG or the VMH, a significant proportion of mice (50%) jumped as many as 35 times over a minute of stimulation (Figures 3H and 3I). In contrast, jumping behavior upon activation of the efferent pathways to either the BNST or the CEA was not significantly different than that observed in eYFP controls (Figures S3F and S3G). Taken together, these finding suggest that the efferent pathways emanating from the dPBN are sufficient to elicit a group of behaviors — running, jumping and analgesia — that would enable escape in the context of injury or other threats (Figure 3I).

Another important component of the response to noxious input is aversion that provides a salient cue to enable avoidance learning. We therefore addressed the degree to which efferent pathways from the lPBN elicit avoidance using a real-time place aversion assay (Figure 4A). As before, experiments were performed on mice in which AAVs encoding either ChR2 or eYFP had been stereotaxically injected into the lPBN and optical fibers were implanted above one of the four efferent targets — CEA, BNST, VMH or lPAG — to enable pathway-selective activation. Notably, regardless of which lPBN efferent pathways that was targeted, ChR2-expressing mice spent significantly less time on the side of the chamber in which they received photostimulation (Figures 4B, 4C, 4D, 4E and 4F).

**Figure 4.**
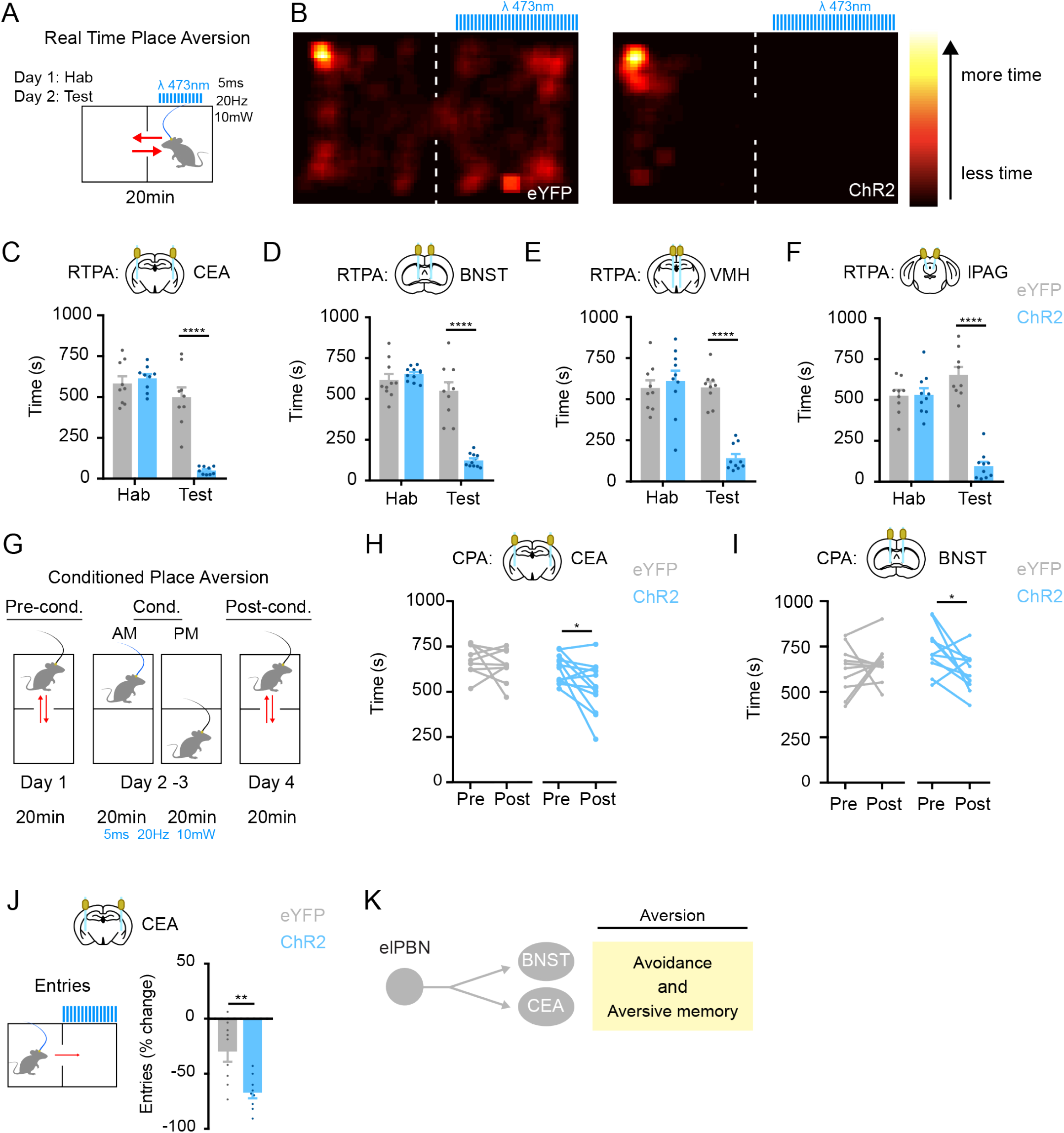
Efferent elPBN projections to BNST and CEA drive aversion. (A) Experimental protocol for real time place aversion (RTPA) assay. Mice were habituated (Hab) for 20 min one day prior to testing (Test). (B) Heat maps of time spent in RTPA chambers upon stimulation of lPBN terminals in CEA in eYFP (left) and ChR2 (right) mice. (C-F) Time spent in photostimulation chamber during habituation phase (Hab) and testing phase (Test) in eYFP (grey) and ChR2 (blue) mice upon stimulation of lPBN efferent terminals in the CEA (C), BNST (D), VMH (E) or lPAG (F). Data are mean ± SEM and dots represent data points from individual animals (n = 9 – 11 mice per group for each experiment). **** Indicates ChR2 mice are significantly different than eYFP controls (Two-way RM ANOVA followed by Holm-Sidak post-hoc test, p < 0.0001). (G) Experimental protocol for conditioned place aversion (CPA). Each session comprises 20 min in the CPA box. (H-I). Photostimulation of elPBN efferent terminals in the CEA (H) or BNST (I) induces CPA. Data are from individual animals (n = 11 - 12 mice per group). * indicates significantly different (Paired Student’s t-test: CEA: p = 0.025; BNST: p = 0.028). (F) Entries into photostimulation chamber upon photostimulation of elPBN efferent terminals in the CEA terminals. ** indicates change in entry number between test phase and habituation phase is significantly different between eYFP and ChR2 mice (unpaired Student’s t-test, p = 0.003). (J) Summary diagram depicting behavioral responses observed upon stimulation of elPBN outputs to BNST and CEA.

Although this behavior was suggestive of aversion, we also considered the possibility that at least in some instances (i.e., VMH and PAG) this apparent avoidance could simply be a consequence of optogenetically-induced locomotion. Thus, to more directly assess whether activation of efferent pathways from the lPBN was sufficient to enable associative conditioning, we used the conditioned place aversion (CPA) assay, in which optogenetic stimulation was selectively paired with one of the two chambers for 20 min on two consecutive days (Figure 4G). When activation of efferent projections to the CEA was the conditioning stimulus, ChR2-expressing mice spent significantly less time on the stimulation-paired side of the chamber (Figure 2H). Similarly, significant CPA was observed when activation of projections to the BNST was used as the conditioning stimulus (Figure 4I). In contrast, repeated photostimulation of efferent projections to either the VMH or the lPAG failed to induce CPA (Figures S3A and S3B). These findings suggest that, although activation of any of the the major outputs from the lPBN results (directly or indirectly) in real time place aversion (RTPA), only those projecting to the CEA or BNST are sufficient for stable aversive learning. To further explore how quickly the mice learned to avoid the side of the chamber in which they receive optogenetic stimulation, we re-analyzed the real time place aversion data, quantifying number of entries into the light-paired chamber. Photostimulation of the efferent projection to the CEA significantly reduced the number of entries (Figure 4J), whereas activation of other efferent projections had no significant effect on entries (Figures S4C, S4D and S4E). These findings reinforce the role of projections to the CEA for avoidance learning because only this cohort of mice showed evidence of learning to avoid the light-paired chamber during the RTPA assay. Together, these data suggest that avoidance memory can be elicited by efferent pathways from the elPBN (Figure 4K), consistent with previous studies (Campos et al., 2018; Chen et al., 2018; Han et al., 2015; Sato et al., 2015).

Having examined the outputs from the lPBN that could mediate the behavioral responses to noxious stimuli, we next characterized the nociceptive inputs to this nucleus. Towards this end, we used the *NK1R^creER^* allele (Huang et al., 2016) to visualize neurokinin 1 receptor-expressing spinoparabrachial neurons, which are known to transmit noxious signals from the spinal cord to the brain (Cameron et al., 2015; Todd, 2010). To visualize the innervation of the lPBN by these neurons, an AAV encoding a Cre dependent fluorescent reporter was injected into the L4-L6 region of the spinal cord of *NK1R^creER^* mice (Figure 5A). We found that *NK1R^creER^* neurons showed dense innervation of the lPBN that was regionally constrained, with the vast majority of these terminals targeting the dPBN and very few targeting the elPBN (Figure 5B), consistent with previous studies (Harrison et al., 2004). To ensure that this observation was not specific to *NK1R^creER^* neurons, we repeated this experiment using a constitutive AAV to label all spinoparabrachial neurons. Again, we saw the same distribution of input from the spinal cord, which was predominant in the dPBN, but not the elPBN (Figures 5C).

**Figure 5.**
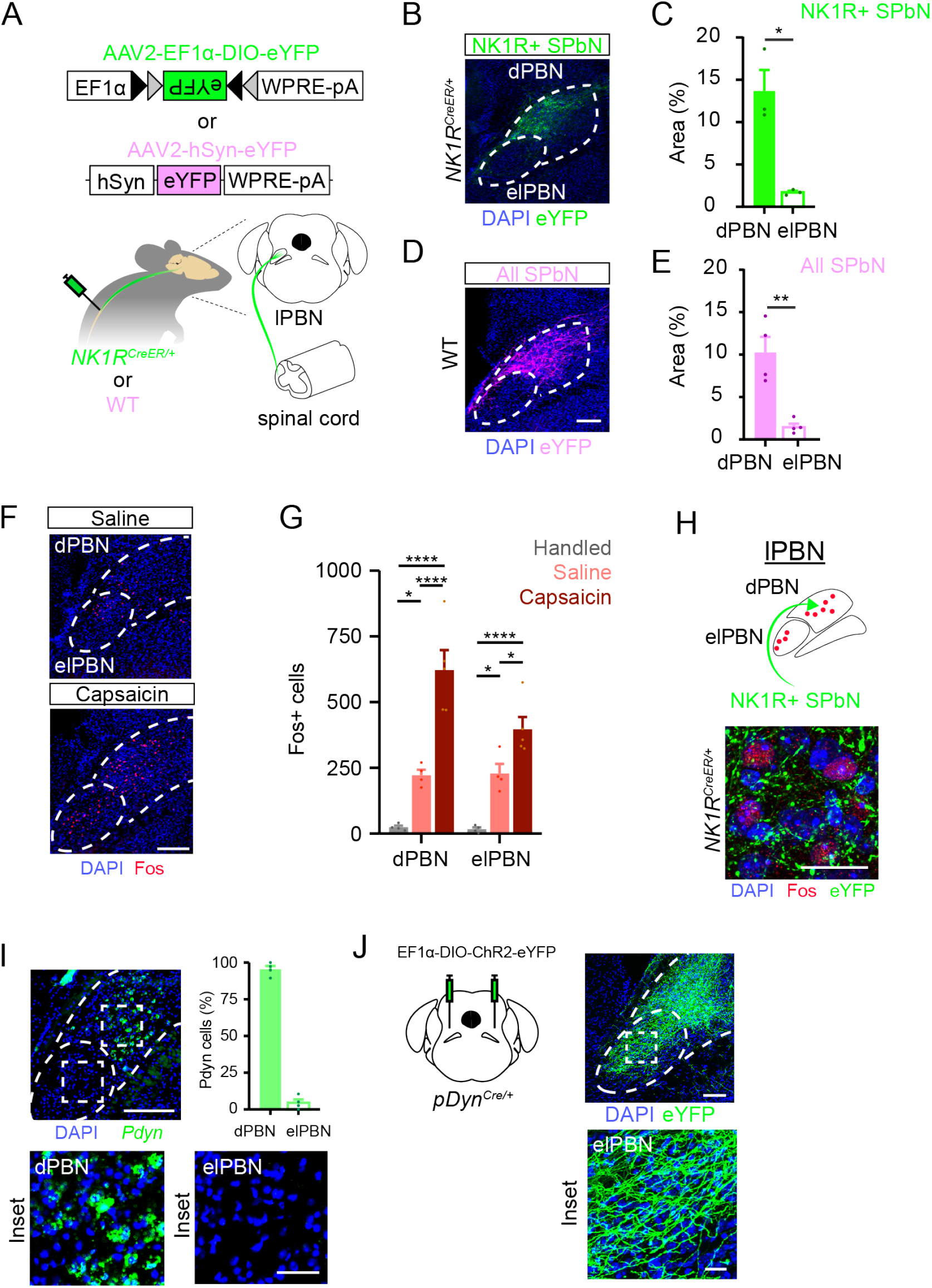
Spinoparabrachial input is concentrated in the dPBN, but noxious stimulation drives Fos expression in both dPBN and elPBN. (A) Experimental strategy to visualize spinal inputs into lPBN. An AAV encoding Cre-dependent eYFP (green) or constitutive eYFP (pseudocolored pink) was injected into the spinal cord (L4-L6) in *NK1R^CreER^* or WT mice. (B-C) Representative image (B) and quantification (C) of the innervation density of efferent terminals in the lPBN from *NK1R^creER^* spinoparabrachial neurons (SPbN). Data are mean ± SEM and dots represent data points from individual animals (n = 3 mice). * indicates area of SPbN projections to dPBN (as percent of region) is significantly greater than that to elPBN (Paired Student’s t-test; p = 0.036). Scale bar = 100 μm. (D-E) Representative image (D) and quantification (E) of the innervation density of efferent terminals in the lPBN from all spinoparabrachial neurons. Data are mean ± SEM and dots represent data points from individual animals (n = 4 mice). * indicates area of spinoparabrachial projections to dPBN (as percent of region) is significantly greater than that to elPBN (Paired Student’s t-test; p = 0.0094). Scale bar = 100 μm. (F-G) Representative image (F) and quantification (G) of Fos induction in the dPBN and elPBN in response to intraplantar saline (10 μl) or capsaicin (10 μl, 0.03%). Data are mean ± SEM and dots represent data points from individual animals (n = 4 - 5 mice per group). Asterisks indicate significantly different (Two-way RM ANOVA followed by Tukey’s post-hoc; * p < 0.05; **** p < 0.001.) Scale bar = 100 μm. (H) *NK1R^creER^* spinoparabrachial terminals (visualized through viral expression of eYFP) are found in close apposition to Fos+ cells in the dPBN following intraplantar capsaicin. Image is representative of data from 4 mice. Scale bar = 25 μm. (I) Representative image and quantification of *pDyn^Cre^* expressing neurons in the dPBN and elPBN as visualized by FISH (n = 4 mice). Scale bar = 100 μm; inset: 25 μm. (J) *pDyn^Cre^* neurons in dPBN project to elPBN. AAV encoding Cre-dependent ChR2 was injected into lPBN of *pDyn^Cre^* mice to visualize projection. Images are representative of data from at least 4 mice. Scale bar = 100 μm; inset: 25 μm.

The paucity of direct nociceptive input to the elPBN was somewhat curious to us in light of previous studies that showed direct innervation of elPBN neurons by spinoparabrachial neurons (Cechetto et al., 1985; Feil and Herbert, 1995; Ma and Peschanski, 1988). Indeed, we found that both the dPBN and elPBN subregions showed significant Fos induction in response to noxious stimulation induced via capsaicin treatment of the hindpaw (Figures 5D and 5E), consistent with previous results (Bernard et al., 1994; Hermanson and Blomqvist, 1996). However, the presynaptic terminals of *NK1R^creER^* spinoparabrachial neurons were only observed in close apposition to Fos+ neurons within the dPBN (Figure 5F).

The apparent discrepancy between the localized nature of the nociceptive input in the dPBN and the widespread nature of the Fos induction by intraplantar capsaicin raised the question of how noxious information reaches the elPBN. With the goal of identifying a neuronal population that might convey nociceptive information between lPBN subregions, we investigated cell types that are known to be expressed in the dPBN using a combination of Cre alleles and stereotaxic injection of Cre dependent AAV reporters to visualize these cells and their projections. In total, six alleles were screened: *SST^cre^*; *NK1R^creER^*, *NTS^cre^*, *CR^cre^*, *CRH^cre^* and *pDyn^cre^* alleles. Although all of these genetic tools uncovered populations of neurons with subregion-specific expression in the lPBN (Figure S5), only the dynorphin population showed localization and anatomy that positioned them to convey noxious information from the dPBN to the elPBN. In particular, using dual fluorescent in situ hybridization (FISH), we found *Pdyn* neurons were located almost exclusively in the dorsal region of the lPBN (Figure 5I), consistent with previous studies (Geerling et al., 2016). Next, we validated the *pDyn^cre^* allele, confirming that Cre-dependent AAV viruses injected into the lPBN of these mice selectively targeted *Pdyn*-expressing neurons (Figure S6A). Finally, using this allele to visualize dynorphin-expressing neurons, we found that dynorphin neurons in dPBN project to the elPBN (Figure 5J). Thus, dynorphin-expressing neurons have cell bodies in the dPBN and send prominent projections to the elPBN.

To further investigate the putative role of dynorphin neurons in nociceptive processing, we used a viral strategy to determine whether spinoparabrachial neurons directly innervate the *pDyn^cre^* subset of dPBN neurons. *pDyn^cre^* mice were stereotaxically injected with AAV encoding a Cre-dependent PSD95-eYFP into the lPBN and another encoding a constitutive synaptophysin-tdtomato into the spinal cord (Figure 6A). These efforts revealed presynaptic terminals from spinal output neurons that were in close apposition to PSD95-eYFP in *pDyn^cre^* neurons, suggestive of direct synaptic contacts. Moreover, we found that intraplantar injection of capsaicin gave rise to strong Fos induction in *pDyn^cre^* neurons. Specifically, 75% of Fos-expressing cells belonged to the *pDyn^cre^* population and Fos was induced in 50% of these cells (Figure 6B). Together, these data provide anatomical and functional evidence that *pDyn^cre^* neurons in the dPBN receive noxious input via spinoparabrachial neurons.

**Figure 6.**
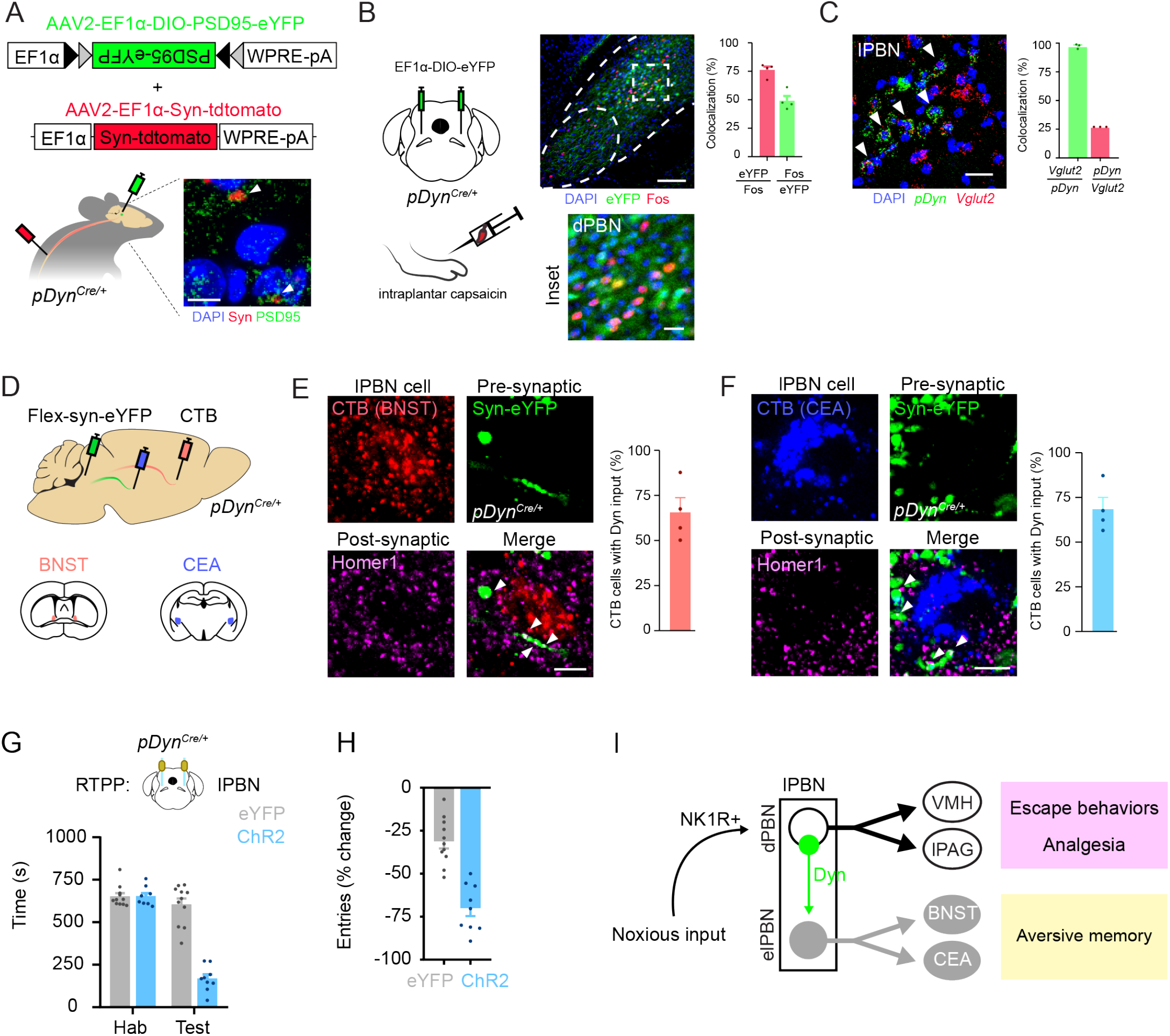
Dynorphin-expressing neurons may convey nociceptive input from dPBN to elPBN. (A) Spinoparabrachial synaptic inputs are found in close apposition to postsynaptic terminals of dynorphin neurons in dPBN. *pDyn^Cre^* mice were injected with AAVs encoding a Cre-dependent PSD95-eYFP in the lPBN and constitutive synaptophysin-tdtomato in the spinal cord (L4-L6). Image is representative of data from 2 mice. (B) *pDyn^Cre^* cells express Fos following in response to noxious stimulation. An AAV encoding Cre-dependent eYFP was stereotactically injected into the lPBN to visualize dynorphin+ cells. Mice received intraplantar injections of capsaicin (10 μl, 0.03%). Representative image and quantification of the colocalization of Fos+ neurons and *pDyn^Cre^* neurons, as visualized by expression of eYFP. Data are mean ± SEM and dots represent data points from individual animals (n = 4 mice). Scale bar = 25 μm; inset: 5 μm. (C) Dynorphin cells are primarily excitatory. Representative image and quantification of colocalization between *pDyn* and *Vglut2* mRNA in dPBN, as observed by dual FISH. Data are mean ± SEM and dots represent data points from individual animals (n = 3 mice). Arrows denote neurons with colocalized signal. Scale bar = 25 μm. (D) Experimental strategy to visualize *pDyn^Cre^* synaptic inputs onto elPBN efferents that project to BNST or CEA. CTB was injected into either the BNST or the CEA and Cre-dependent synaptophysin-eYFP was delivered into the lPBN of *pDyn^Cre^* mice. (E-F). Synaptic terminals from *pDyn^Cre^* neurons (green) are in close apposition to Homer1 puncta (purple) surrounding CTB-labeled neurons that have been backlabeled from the BNST (E; red) or CEA (F; blue). Data are mean ± SEM and dots represent data points from individual animals (n = 4 mice per experiment). (G) Photostimulation of ChR2-expressing *pDyn^Cre^* cells in the dPBN elicits real time place aversion. Data are mean ± SEM and dots represent data points from individual animals (n = 9 – 11 mice per group). **** indicates significantly different (Two-way RM ANOVA followed by Holm-Sidak post-hoc test, p < 0.0001). (H) Photostimulation of ChR2-expressing *pDyn^Cre^* cells in the dPBN significantly diminished entry number into the stimulation chamber. Data are mean ± SEM and dots represent data points from individual animals (n = 9 – 11 mice per group). **** indicates significantly different (unpaired Student’s t-test p < 0.0001). (I) Model: Noxious input is conveyed primarily to the dPBN. Efferents from the dPBN collateralize to VMH and lPAG and mediate behavioral responses that enable escape. Dynorphin neurons in the dPBN convey noxious information to elPBN. Efferents from the elPBN collateralize onto the CEA and BNST and mediate aversion and avoidance memory.

To characterize these *pDyn^cre^* neurons in more detail, we next examined whether this population represented an excitatory or inhibitory population of neurons through dual fluorescent in situ hybridization (FISH). We found that nearly all *Pdyn* transcript colocalized with *Vglut2*, with *Pdyn* cells representing approximately one-quarter of the excitatory population within the dPBN (Figure 6C). In contrast, there was very little to no overlap of *Pdyn* and the inhibitory marker *Vgat* (Figure S6B). Thus, from a neurochemical standpoint, dynorphin neurons in the dPBN are positioned to relay nociceptive information to the elPBN.

To further investigate whether dynorphin neurons could provide a cellular substrate for transmission of nociceptive information to elPBN efferents, we used viral and retrograde tracing approaches to determine whether *pDyn^cre^* neurons form anatomical connections with elPBN neurons that project to CEA and BNST. Towards this end, we stereotaxically injected AAV encoding a Cre-dependent synaptophysin-eYFP into dPBN together with CTB into either the CEA or BNST of *pDyn^cre^* mice (Figure 6D). These experiments suggested that approximately two-thirds of CTB-labeled cells from either the BNST or the CEA receive synaptic input from *pDyn^cre^* neurons as supported by the close apposition of retrogradely labeled cells to synaptophysin-eYFP and the post-synaptic density marker Homer1 (Figures 6E and 6F). These data provide anatomical support for the idea that *pDyn^cre^* neurons may convey noxious information to neurons in the elPBN that have efferent projections to the CEA and/or BNST.

To further test the idea that *pDyn^cre^* neurons in the dPBN convey information to the elPBN, we next examined whether activation of this population is sufficient to elicit the behavioral responses that are mediated by elPBN efferents. To manipulate these cells, we delivered an AAV encoding Cre-dependent ChR2 or eYFP into the lPBN of *pDyn^cre^* mice. Consistent with our hypothesis, we found that photostimulation of *pDyn^cre^* neurons in the lPBN mice gave rise to aversive behaviors, but not escape behaviors. In particular, optogenetic stimulation resulted in real time place aversion coupled with a significant reduction in number of entries into the stimulation chamber (Figures 6G and 6H). In contrast, the activation of *pDyn^cre^* neurons had no effect on escape behaviors including running, jumping or tail flick latency (Figures S6C, S6D and S6E). These data highlight that *pDyn^cre^* neurons serve a crucial link for the recruitment of elPBN pathways to CEA and BNST.

## Discussion

We have identified two anatomically and functionally distinct populations of lPBN neurons that underlie different aspects of the nocifensive response. In dPBN, neurons receive direct input from spinal projection neurons and mediate behaviors that would enable escape. In elPBN, neurons are activated by a neurochemically defined population of dPBN neurons that express dynorphin and mediate aversive learning. Together, these findings provide evidence for a neural substrate that coordinates diverse behavioral responses to noxious stimuli (Figure 6I).

Although this study explores the idea that different efferent pathways from the lPBN mediate different components of the pain response, we acknowledge that such a model is undoubtedly an oversimplification of the complex and interconnected circuitry underlying these behaviors. The optogenetic approach used herein provides a new level of specificity by directly activating lPBN efferents at a given target. However, multiple brain regions are likely to have been affected indirectly as a consequence of this manipulation. For instance, efferents from the BNST project back to the lPBN, as well as to the CEA, the lPAG, the RVM, and numerous other brain structures (Barik et al., 2018; Rodriguez et al., 2017; Tokita et al., 2009). Consistent with this interconnectivity, we found that photostimulation of a single efferent pathway activates multiple brain regions, as evidenced by Fos induction. It is therefore not surprising that some behaviors were not completely specific to the activation of a particular lPBN projection. Most notably, stimulation of efferents to the BNST elicited small but significant escape responses (running and analgesia), in addition to aversive learning. Nevertheless, the broad findings of our study are consistent with a modular output from the lPBN that would enable the coordination of nocifensive responses in a context-dependent manner.

It is intriguing that distinct lPBN efferents would be predicted to have opposite effects on the pain experience: those emanating from the dorsal division would be expected to decrease pain, while those from the external lateral domain would be expected to exacerbate pain. The efferent pathway from the dPBN might predominate in the context of an emergency to help avoid injury, whereas the efferent pathway from the elPBN might predominate once imminent danger has past to facilitate aversive learning. The neural substrate for the coordination of different efferent responses in this way is poorly understood. Our work suggests that *pDyn^Cre^* neurons may be involved in this coordinated regulation between efferent projections emanating from the dorsal and external lateral domains, respectively. Our data reveal that *pDyn^Cre^* neurons have cell bodies in the dPBN but send extensive projections to the elPBN and, consistent with this anatomy, we find that these cells are activated by noxious input and drive aversion, but not escape behaviors. However, we note that this is unlikely to be the only function of *pDyn^Cre^* cells in the lPBN because these neurons have been shown to play important roles in temperature homeostasis (Geerling et al., 2016; Nakamura and Morrison, 2008, 2010). These findings raise the possibility that *pDyn^Cre^* neurons in the lPBN are not a single, homogeneous population. In future studies, it will be important to characterize this heterogeneity in more detail to identify bona fide cell types and characterize how each responds to diverse stimuli.

More than any other species, humans have a detailed cortical representation that informs conscious perception of pain. But this cortico-centric view of pain may overlook the fundamental idea that avoiding tissue damage is a primal need for which subcortical pathways play a central role. Our studies highlight a potentially important role of dynorphin in the lPBN in this regulation. Because chronic pain has such a profound effect on mental health and well-being, further studies investigating changes in this circuitry in the context of chronic pain and the possible role of dynorphin signaling therein are warranted.

## STAR METHODS

### Animals

Mice were given free access to food and water and housed under standard laboratory conditions. The use of animals was approved by the Institutional Animal Care and Use Committee of the University of Pittsburgh. *Pdyn-IRES-Cre* (Krashes et al., 2014), *Gad2-IRES-Cre* (Taniguchi et al., 2011), *NK1R-CreER* (Huang et al., 2016), *Ai34(RCL-Syp/tdT)-D*, and *Rosa26 CAG-LSL-ReaChR-mCit* (Hooks et al., 2015) were obtained from Jackson Laboratory. Wild-type C57BL/6 mice were obtained from Charles River (Cat # 027). For all experiments 8 – 16 week-old male and female mice were used. In all cases, no differences between male and female mice were observed and so the data were pooled. Age-matched littermates were used for all behavioral experiments that involved mice harboring the knock-in allele Cre-recombinase.

### Viruses

The following viruses were used for experimentation: AAV2-hsyn-eYFP (Addgene: 50465), AAV2-hSyn-hChR2(H134R)-eYFP (Addgene: 26973), AAV2-EF1a-DIO-eYFP (Addgene: 27056), AAV2-EF1a-DIO-hChR2(H134R)-eYFP (Addgene: 20298), AAV9-CAGGS-FLEX-ChR2-tdtomato.WRPE.SV40 (Addgene: 18917), AAV8.2-hEF1a-DIO-synaptophysin-eYFP (MGH: AAV-RN2), AAV8.2-hEF1a-DIO-PSD95-eYFP (MGH: AAV-RN7), and AAV8.2-hEF1a-synaptophysin-mCherry (MGH: AAV-RN8). Viruses were purchased from University of North Carolina Vector Core, University of Pennsylvania Vector Core, and Massachusetts Gene Technology Core.

### Stereotaxic injections and implantation of optical fiber

Animals were anesthetized with 2% isoflurane and placed in a stereotaxic head frame. Ophthalmic ointment was applied to the eyes. The scalp was shaved, local antiseptic applied (betadine), and a midline incision made to expose the cranium. The skull was aligned using cranial fissures. A drill bit (MA Ford, #87) was used to create a burr hole and custom made metal needle (33 gauge) loaded with virus was subsequently inserted through the hole to the injection site. Virus was infused at a rate of 100nL/min using a Hamilton syringe with a microsyringe pump (World Precision Instruments). Wildtype mice received 0.150 μl of virus. All other Cre-expressing mice received 0.5 μl virus. The injection needle was left in place for an additional 5-10 min and then slowly withdrawn. Injections and optical fiber implantations were performed bilaterally at the following coordinates for each brain region: BNST: AP +0.50 mm, ML ± 1.00 mm, DV −4.30; CEA: AP −1.20 mm, ML ± 2.85 mm, DV −4.50; VMH: AP −1.48 mm ML ± −0.50 mm DV −5.80 mm; lPAG: AP −4.70 mm, ML ± 0.74 mm, DV: −2.75; and lPBN AP −5.11 mm, ML ± 1.25 mm, DV: −3.25. For implantation of optical fibers (Thor Labs: 1.25 mm ceramic ferrule 230 μm diameter), implants were slowly lowered 0.300-0.500 mm above the site of injection and secured to the skull with a thin layer of Vetbond (3M) and dental cement. The incision was closed using Vetbond and animals were given a subcutaneous injection of buprenorphine (0.3mg/kg) and allowed to recover over a heat pad. Mice were given 4 weeks to recover prior to experimentation.

### RNAscope in situ hybridization

Multiplex fluorescent in situ hybridization was performed according to the manufacturer’s instructions (Advanced Cell Diagnostics #320850). Briefly, 18 μm-thick fresh-frozen sections containing the parabrachial nucleus were fixed in 4% paraformaldehyde, dehydrated, treated with protease for 15 minutes, and hybridized with gene-specific probes to mouse *Pdyn* (#318771), *Slc32a1* (#319191), and *Slc17a6* (#319171). DAPI (#320858) was used to visualize nuclei. 3-plex positive (#320881) and negative (#320871) control probes were tested. Two to three 18 μm z-stacked sections were quantified for a given mouse, and 2 – 4 mice were used per experiment.

### Immunohistochemistry

Mice were anesthetized with an intraperitoneal injection of urethane, transcardially perfused, and post-fixed at least four hours in 4% paraformaldehyde. 40 or 65 μm thick transverse brain or spinal cord sections were collected on a vibratome and processed free-floating for immunohistochemistry. Sections were blocked at room temperature for two hours in a 10% donkey serum, 0.1% triton, 0.3M NaCl in phosphate buffered saline. Primary antisera was incubated for 14 hours overnight at 4°C (except for rabbit anti-Homer1, detailed below): rabbit anti-c-Fos (1:5K), mouse NeuN (1:1K), chicken anti-GFP (1:1K), rabbit anti-NK1R (1:1K), and rabbit anti-Homer1 (1:1K, incubated for 3 days). Sections were subsequently washed three times for 20 minutes in wash buffer (1% donkey serum, 0.1% triton, 0.3M NaCl) and incubated in secondary antibodies (Life Technologies, 1:500) at room temperature for two hours. Sections were then incubated in Hoechst (ThermoFisher, 1:10K) for 1 minute and washed 7 times for 15 minutes in wash buffer, mounted and coverslipped.

### CTB backlabeling

Fluorescently conjugated cholera toxin subunit B-Alexa-fluor conjugates −555 and −647 (CTB, ThermoFisher C34778, C22843) were stereotactically injected (0.2 μl, 1mg/ml) into the brain regions of interest and subsequently analyzed 10 days following injection. Mice were perfused and brains were processed as described above for immunohistochemistry. CTB-labeled cells were quantified using 65 μm z-stacked images at 2 μm steps of the entire lPBN (n = 3 – 5 mice per backlabeled region). For retrograde labeling of cells and quantification of pre- and post-synaptic markers, 3 – 4 40 μm sections were quantified for a given animal, and 4 mice were used per experiment.

### Image acquisition and quantification

Full-tissue thickness sections were imaged using either an Olympus BX53 fluorescent microscope with UPlanSApo 4x, 10x, or 20x objectives or a Nikon A1R confocal microscope with 20X or 60X objectives. All images were quantified and analyzed using ImageJ. For all images, background pixel intensity was subtracted as calculated from control mice. To quantify the area of synapses observed, confocal images using single optical planes were converted into a binary scale and area of signal taken as a ratio of the total area (one section per region of interest, n = 6 mice). To quantify CTB-labeled cells in tracing experiments, confocal images were manually quantified using full-tissue thickness z-stacked images at 2 μm steps of the entire lPBN (3 – 4 mice per group). To quantify images in RNAscope in situ hybridization experiments, confocal images of tissue samples (1 – 2 sections per mouse over 2 – 4 mice) were imagined and only cells whose nuclei were clearly visible by DAPI staining and exhibited fluorescent signal were counted. To quantify Fos-labeled cells, 65 μm sections of the entire lPBN were imaged using the fluorescent microscope and images manually counted.

### Fos induction (intraplantar capsaicin)

Fos induction was performed as previously described in (Rodriguez et al., 2017). Mice were lightly anesthetized with isoflurane and received one of the following treatments: handled (no injection), 10 μl unilateral intraplantar saline, or 10 μl unilateral intraplantar capsaicin (0.03% capsaicin w/v in 2.5% Tween 80 and 2.5% ethanol in PBS). Mice were then placed back into their cages and subsequently perfused 90 minutes later and neural tissue collected according to protocol for immunohistochemistry.

### Behavior

All assays were performed and scored by an experimenter blind to virus (eYFP or ChR2). Post hoc analysis confirming specificity of viral injections and proper fiber implantation were also performed blinded to animal identity, and mice in which viral injections and/or fiber implantation were considered off target excluded from analysis. All testing was performed in the University of Pittsburgh Rodent Behavior Analysis Core. Optogenetic stimulation parameters were determined empirically as follows: 10mW, 20Hz, 5ms duration pulses.

### Real time place aversion assay (RTPA)

Mice were stereotaxically injected with either channelrhodopsin or control eYFP virus and optical fibers implanted at the downstream terminals of interest. Four weeks following injection mice were habituated to a custom-made 2-chamber (40cm × 28cm × 20cm chamber) for real time place aversion testing. Mice were habituated on day 1 for 20 minutes and subsequently tested the next day for 20 minutes. Light stimulation was delivered whenever the mouse entered one of two sides of the chamber and turned off when the animal exited that chamber. The side of stimulation was counterbalanced. The behavior was recorded and post-hoc analysis performed to determine body position using the open source software Optimouse (Ben-Shaul, 2017). Position data were discarded according to established criteria, and velocity was computed as described here (https://www.biorxiv.org/content/10.1101/558643v1).

### Tail immersion test

Mice were habituated to mice restraints 15 minutes for 5 days before testing. Tails were immersed 3 cm into a water bath at 48°C or 55°C, and the latency to tail flick was measured three times per temperature with a 1 minute interval between trials. For optogenetic testing, mice were photostimulated for 10 seconds prior to tail immersion testing.

### Escape response test

Mice were placed in an open field chamber and allowed to habituate for five minutes before two 30-second optogenetic stimulation bouts and two minute resting period between bouts. The behavior was recorded and post-hoc analysis performed to determine body position using the open source software Optimouse as described in RTPA.

### Conditioned place aversion test

Mice were placed in a two-chamber plexiglass box for 20 minutes and allowed to freely roam between one of two sides differentiated by visual cues (spots versus stripes). For two conditioning days, mice were restricted to one of two sides and received either no stimulation or photostimulation (3 seconds on, 2 seconds off at 20Hz, 5ms pulse duration, 10mW) for 20 minute periods in the morning and afternoon. On the test day, mice were placed back into the box and allowed to freely explore either chamber. The behavior was recorded and post-hoc analysis performed to determine body position using the open source software Optimouse as described in RTPA.

### Mechanical allodynia

Mice were allowed to habituate for at least two hours prior to testing. Mice received a 10ul intraplantar injection of 0.03% capsaicin dissolved in 2.5% Tween, 2.5% ethanol in PBS and tested for mechanical hypersensitivity via the up-down method (Chaplan et al., 1994). After 5-10 minute resting period, mice were optogenetically stimulated and tested for mechanical hypersensitivity. Mice were again allowed to rest for 5-10 minutes before von Frey testing for post-stimulation effects on mechanical hypersensitivity.

### Intraspinal injections

Mice were anesthetized with 100mg/kg ketamine and 10 mg/kg xylazine. An incision was made at the spinal cord level corresponding to L4-6 dermatome. The intrathecal space was exposed, and two injections of approximately 1μl of virus was infused 300 μm below the surface of the spinal cord at 100nL/min via glass pipette through the intrathecal space corresponding to L4-6 of the spinal cord. The glass pipette was left in place for an additional 5 minutes before withdrawal. The incision was closed with 5-0 vicryl suture. Buprenorphine was delivered post-surgery at 0.3mg/kg subcutaneously, and mice were allowed to recover over a heat pad.

### Statistical analyses

All statistical analyses were performed using GraphPad Prism 7.0. Values are presented as mean ± SEM. Statistical significance was assessed using Fisher’s exact test for categorical data, Students t-test, or two-way repeated measures ANOVA followed by Holm-Sidak post-hoc test. Significance was indicated by p ≤ 0.05. Sample sizes were based on pilot data and are similar to those typically used in the field.

#### Data availability

The data collected in this study are available from the corresponding author upon request.

#### Code availability

All custom-written MatLab code used in this study is available upon request.

## ACKNOWLEDGEMENTS

We thank Hannah E. Piston for generating equipment for tail flick assay and Michael S. Gold for helpful comments. Research reported in this publication was supported by the Virginia Kaufman Endowment Fund, NIH/NIAMS grant AR063772, NIH/NINDS grant NS096705 (S.E.R.), NRSA F30 grant F30NS096860 and NIGM/NIH T32GM008208 (M.C.C).

## AUTHOR CONTRIBUTIONS

Conceptualization: M.C.C and S.E.R.

Methodology: M.C.C, E.K.N, A.E.P and S.E.R.

Investigation: M.C.C and S.E.R.

Writing: M.C.C and S.E.R.

Funding Acquisition: M.C.C and S.E.R.

## CONTACT FOR REAGENT AND RESOURCE SHARING

Further information and requests for resources and reagents should be directed to and will be fulfilled by the corresponding authors and Lead Contacts, Dr. Sarah E. Ross.

## KEY RESOURCES TABLE

| REAGENT or RESOURCE | SOURCE | IDENTIFIER |
| --- | --- | --- |
| **Antibodies** |  |  |
| Rabbit anti-homer 1 | Frontier Institute | Cat: AB_2571774; RRID: AB_2571774 |
| Chicken anti-GFP | Aves Laboratory | Cat: GFP-1020; RRID: AB_10000240 |
| Rabbit anti-c-fos | Santa Cruz Biotech | Cat: Sc-52; RRID: AB_216783 |
| Mouse anti-NeuN | Millipore | Cat: MAB377; RRID: AB_2298772 |
| Rabbit anti-NK1R | Sigma Aldrich | Cat: SAB4502913; RRID: AB_10746598 |
| Donkey anti-chicken (IgG) Alexa Fluor 488 secondary antibody | Jackson ImmunoResearch | Cat: 703-035-155; RRID: AB_2340375 |
| Donkey anti-rabbit (IgG) Alexa Fluor 555 secondary antibody | ThermoFisher | Cat: A-31572; RRID: AB_162543 |
| Donkey anti-mouse (IgG) Alexa Fluor 647 | ThermoFisher | Cat: A-31571; RRID: AB_162542 |
| **Bacterial and Virus Strains** |  |  |
| AAV2-hSyn-eYFP | UNC | Addgene: 50465 |
| AAV2-hSyn.hChr2(H134R).eYFP | UNC | Addgene: 26973 |
| AAV2-EF1a-DIO-eYFP | UNC | Addgene: 27056 |
| AAV2-EF1a-DIO-hChr2(H134R)-eYFP | UNC | Addgene: 20298 |
| AAV8.2-hEF1a-DIO-synaptophysin-eYFP | MGH GTC | AAV-RN2 |
| AAV8.2-hEF1a-DIO-PSD95-eYFP | MGH GTC | AAV-RN7 |
| AAV8.2-hEF1a-synaptophysin-mCherry | MGH GTC | AAV-RN8 |
| AAV9-CAGGS-FLEX-ChR2-tdtomato.WRPE.SV40 | Penn Vector Core | Addgene: 18917 |
| **Chemicals, Peptides, and Recombinant Proteins** |  |  |
| Cholera toxin subunit B (Recombinant), Alexa Fluor 555 | ThermoFisher | C34778 |
| Cholera toxin subunit B (Recombinant), Alexa Fluor 647 | ThermoFisher | C22843 |
| PFA | Sigma Aldrich | P6148 |
| Capsaicin | Sigma Aldrich | M2028 |
| **Critical Commercial Assays** |  |  |
| **RNAscope** |  |  |
| Fluorescent multiplex assay | ACD | 320850 |
| *Pdyn* probe | ACD | 318771 |
| *Slc32a1* probe | ACD | 319191 |
| *Slc17a6* probe | ACD | 319171 |
| EYFP probe | ACD | 312131 |
| DAPI | ACD | 320858 |
| 3-plex positive control probe | ACD | 320881 |
| 3-plex negative control probe | ACD | 320871 |
| **Experimental Models: Organisms/Strains** |  |  |
| Mouse: C57BL6 | Charles River | 027 |
| Mouse: Pdyn-IRES-Cre<br>B6;129S-Pdyn<sup>tm1.1(cre)Mjkr</sup>/LowlJ | Jackson Laboratory<br>Krashes et al. 2014 | IMSR Cat:<br>JAX:027958; RRID:<br>IMSR_JAX:027958 |
| Mouse: Gad2-IRES-Cre<br>Gad2<sup>tm2(cre)Zjh</sup>/J | Jackson Laboratory<br>Taniguchi et al. 2011 | IMSR Cat:<br>JAX:010802; RRID:<br>IMSR_JAX:010802 |
| Mouse: NK1R<sup>CreER</sup> | Ross lab; Huang et al.<br>2016 | N/A |
| Mouse: Ai34D or Ai34(RCL-Syp/tdT)-D<br>B6;129S-Gt(ROSA)26Sor<sup>tm34.1(CAG-Syp/tdTomato)Hze</sup>/J | Jackson Laboratory<br>Zeng 2011 | IMSR Cat:<br>JAX:012570; RRID:<br>IMSR_JAX:012570 |
| Mouse: Rosa26 CAG-LSL-ReaChR-mCit<br>B6.Cg-Gt(ROSA)26Sor<sup>tm2.2Ksvo</sup>/J | Jackson Laboratory<br>Hooks et al. 2015 | IMSR Cat:<br>JAX:026294; RRID:<br>IMSR_JAX:026294 |
| Software and Algorithms |  |  |
| Prism 7.0 | GraphPad | N/A |
| Matlab | Mathworks | N/A |
| Mouse tracking algorithm | https://www.biorxiv.org<br>/content/10.1101/5586<br>43v1 | N/A |

## Supplemental Information

### Supplemental Data

**Figure S1.**
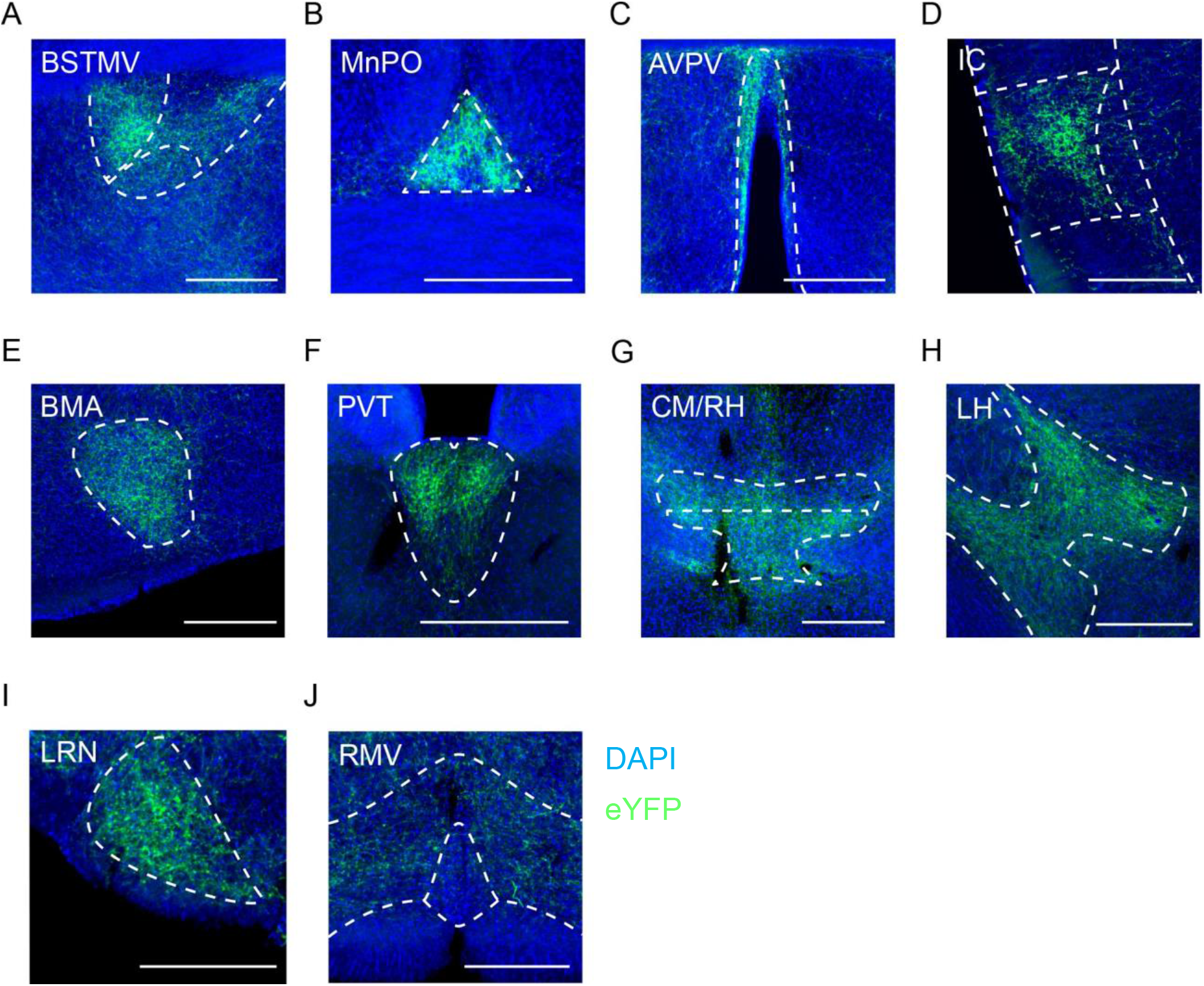
Efferents from the lPBN target numerous brain regions. Efferents from the lPBN targets numerous brain regions, as visualized following stereotaxic injection of an AAV encoding eYFP into the lPBN (0.2 μl, bilateral). **(A)** Medial ventral division of bed nucleus stria terminalis (BNSTMV). **(B)** Median preoptic nucleus (MnPO). **(C)** Anteroventral periventricular nucleus (AVPV). **(D)** Insular cortex. **(E)** Basomedial amygdaloid nucleus (BMA). **(F)** Paraventricular thalamic nucleus (PV). **(G)** Centromedial thalamic nucleus (CM) and rhomboid nucleus (RH). **(H)** Lateral hypothalamus (LH). **(I)** Lateral reticular nucleus (LRN). **(J)** Rostral ventromedial medulla (RVM). Scale bar = 200 μm (A - J)

**Figure S2.**
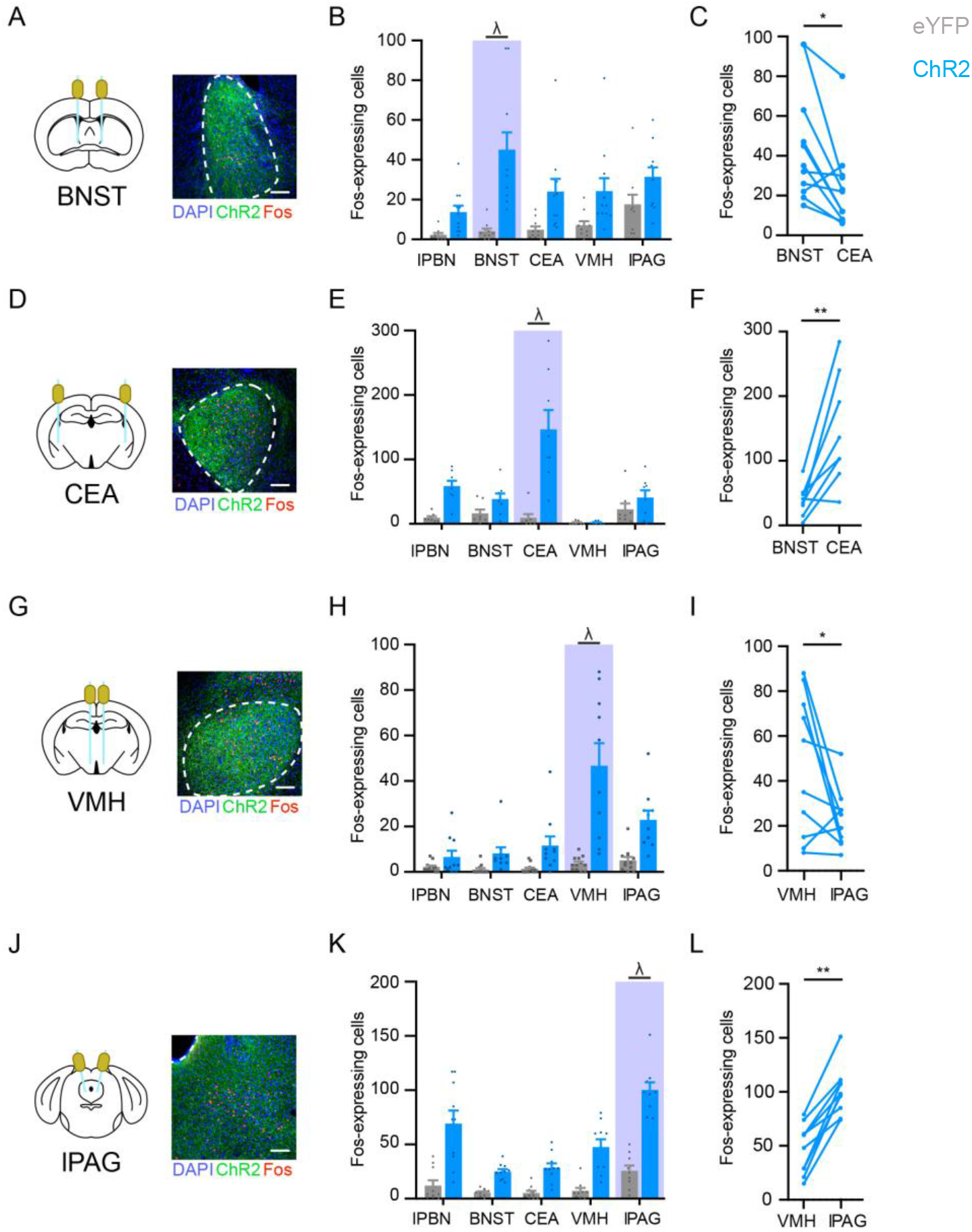
Pathway selective photostimulation revealed through Opto-Fos experiments. To assess the degree to which photstimulation of a specific lPBN efferent was selective, Fos induction was analyzed 90 min after photostimulation (20 min; 20 Hz, 5 ms pulses, 10 mW). **(A)** Diagram and image of ChR2-expressing terminals and Fos-expressing cells in BNST. **(B)** Fos induction in eYFP-expressing mice (control, grey) and ChR2-expressing mice (blue) upon photostimulation of lPBN efferent terminals in the BNST. Data are mean ± SEM and dots represent data points from individual animals (n = 8 - 11 mice per group). **(C)** The number of Fos-expressing cells in BNST is significantly greater than that in CEA (paired Student’s t-test p = 0.01, n = 11 mice). **(D)** Diagram and image of ChR2-expressing terminals and Fos-expressing cells in CEA. **(E)** Fos induction in eYFP-expressing mice (control, grey) and ChR2-expressing mice (blue) upon photostimulation of lPBN efferent terminals in the CEA. Data are mean ± SEM and dots represent data points from individual animals (n = 8 - 9 mice per group) **(F)** The number of Fos-expressing cells in CEA is significantly greater than that in BNST (paired Student’s t-test p = 0.004, n = 9 mice). **(G)** Diagram and image of ChR2-expressing terminals and Fos-expressing cells in VMH. **(H)** Fos induction in eYFP-expressing mice (control, grey) and ChR2-expressing mice (blue) upon photostimulation of lPBN efferent terminals in the VMH. Data are mean ± SEM and dots represent data points from individual animals (n = 10 - 11 mice per group). (I) The number of Fos-expressing cells in VMH is significantly greater than that in lPAG (paired Student’s t-test, p = 0.04, n = 10). **(J)** Diagram and image of ChR2-expressing terminals and Fos-expressing cells in VMH. **(K)** Fos induction in eYFP-expressing mice (control, grey) and ChR2-expressing mice (blue) upon photostimulation lPBN efferent terminals in the VMH. Data are mean ± SEM and dots represent data points from individual animals (n = 9 - 10 mice per group) **(L)** The number of Fos-expressing cells in lPAG is significantly greater than that in VMH (paired Student’s t-test p = 0.004, n = 9). Scale bar = 100 μm (A, D, G, J)

**Figure S3.**
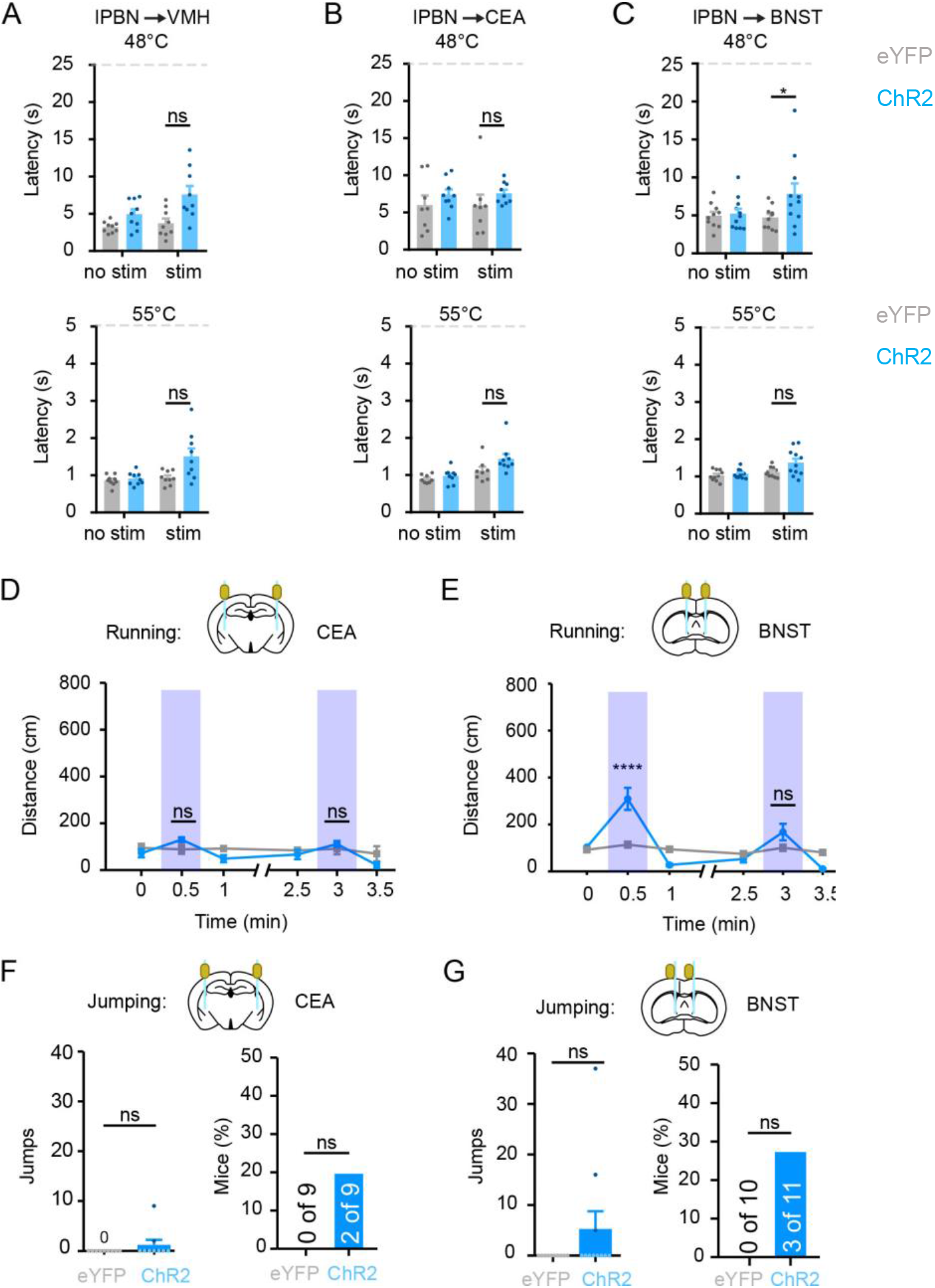
Effect of photoactivation of lPBN efferents on descending modulation, running, and jumping. **(A-B)** Optogenetic activation of terminals in VMH or CEA of ChR2-injected mice (blue bars) does not increase latency of tail flick response to low or high noxious heat compared to eYFP-expressing controls (grey). Data are mean ± SEM and dots represent data points from individual animals (n = 8 - 9 mice per group; ns, not significant p > 0.05, two-way RM ANOVA). **(C)** Photostimulation of projections to BNST in ChR2-expressing mice (blue bars) increases latency to tail flick at 48 °C but not 55 °C compared to control mice (grey bars). Data are mean ± SEM and dots represent data points from individual animals (n = 10 - 11 mice per group; * indicates significant, two-way RM ANOVA followed by Holm-Sidak post-hoc test, p = 0.03). **(D)** Optogenetic stimulation of terminals in CEA of ChR2-injected mice has no effect on lateral movement. Data are mean ± SEM (n = 8 – 9 mice per group; ns, not significant, two-way RM ANOVA, p > 0.05). **(E)** Optogenetic stimulation of terminals in BNST promotes significant lateral movement in the first but not second bout of photostimulation. Data are mean ± SEM (n = 10 - 11 mice per group, **** indicates significant difference, two-way RM ANOVA followed by Holm-Sidak post-hoc test, p < 0.0001). **(F - G)** The number of jumps or proportion of mice exhibiting jumping was not significantly higher in ChR2-expressing mice that were stimulated in CEA or BNST compared to eYFP controls (Mann-Whitney t-test: CEA: p = 0.47; BNST: p = 0.21, Fisher’s exact test: CEA: p = 0.47; BNST: p = 0.21). Data are mean ± SEM and dots represent data points from individual animals (n = 9 - 11 mice per group).

**Figure S4.**
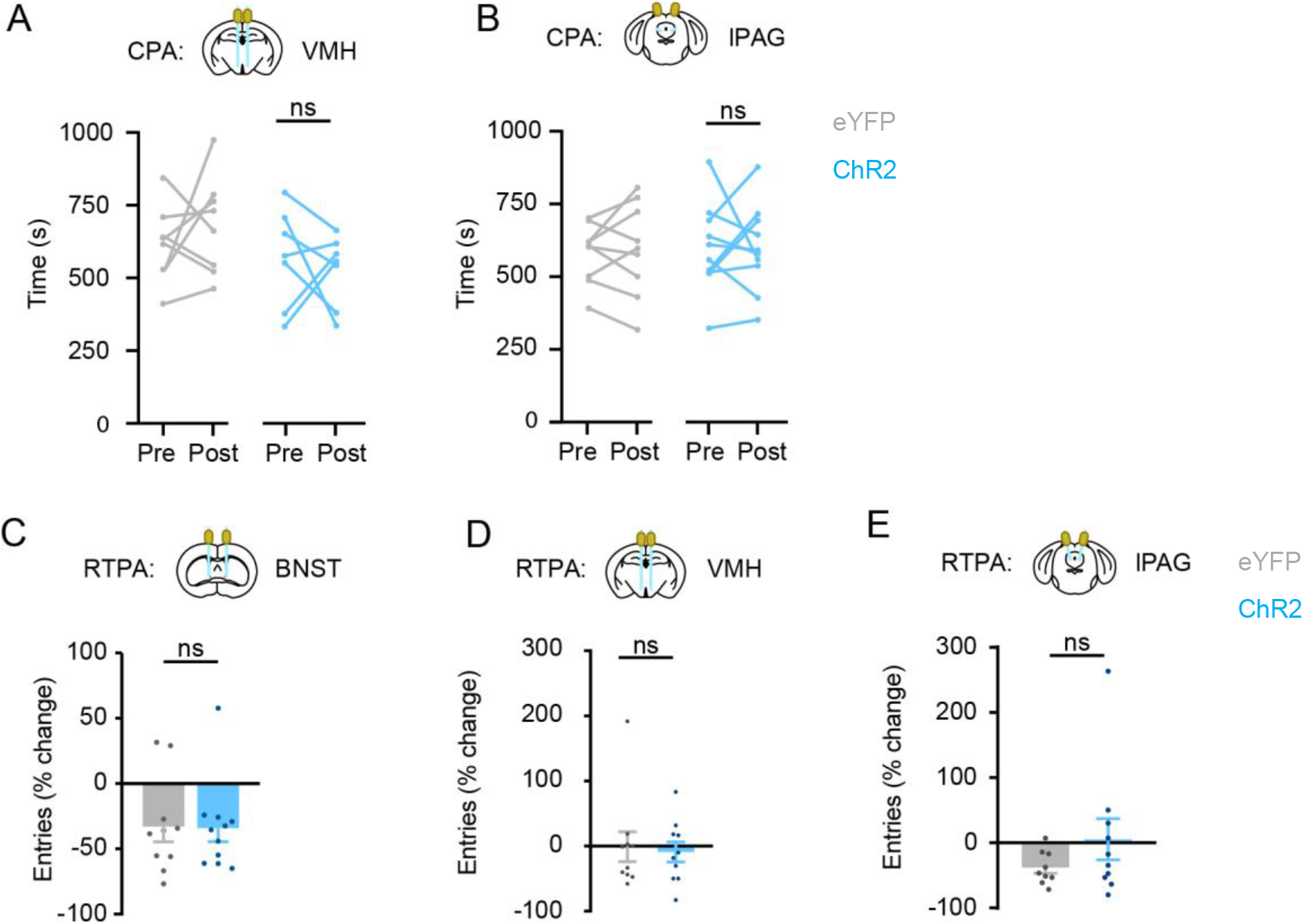
Optogenetic stimulation of lPBN efferents and aversive memory. **(A-B)** Optogenetic activation of terminals in VMH (A) or lPAG (B) of ChR2-injected mice does not cause CPA (paired Student’s t-test: VMH: p = 0.605; lPAG: p = 0.986). Data are from individual mice (n = 7 – 9 mice per condition). **(C-E)** Optogenetic activation of ChR2-expressing efferents terminals in BNST (C), VMH (D), or lPAG (E) does not affect the number of entries into the photostimulation chamber during RTPA (Student’s t-test, BNST: p = 0.942; VMH: p = 0.772; lPAG: p = 0.219). Data are mean ± SEM and dots represent data points from individual animals (n = 9 - 11 mice per group).

**Figure S5.**
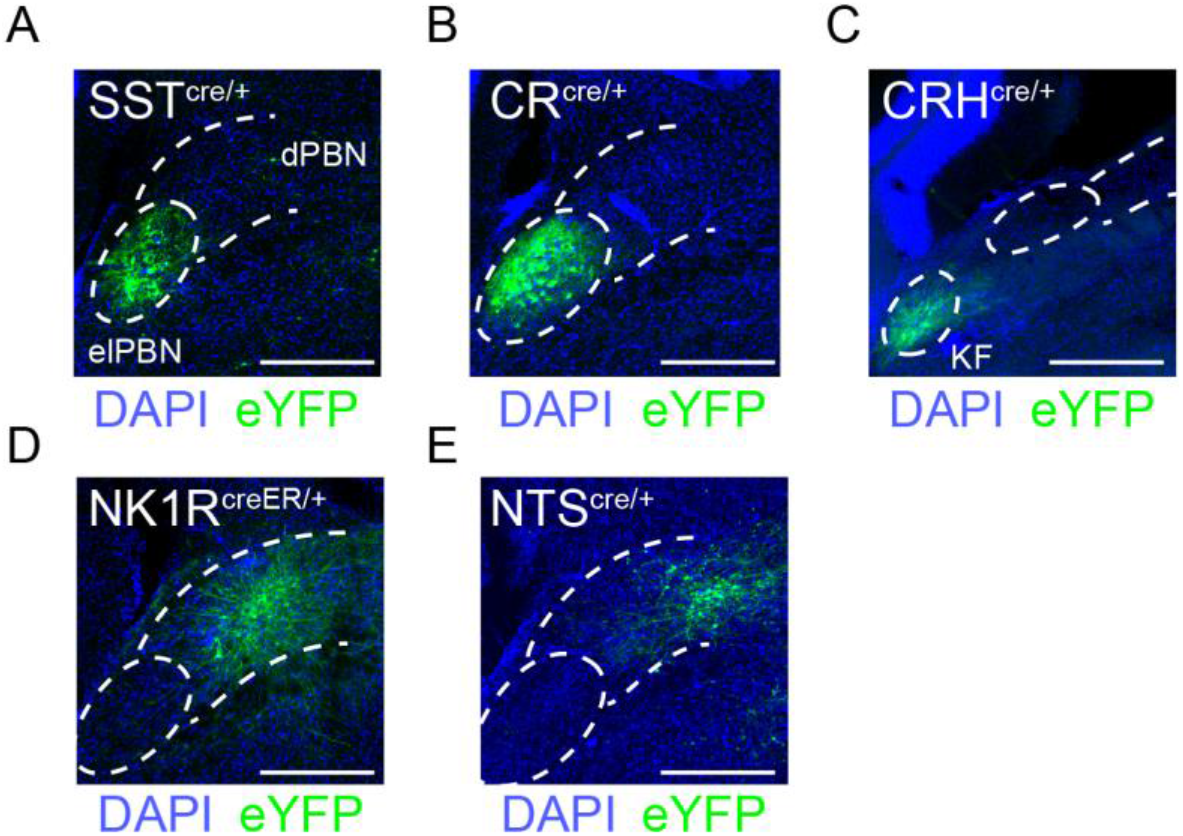
Screen of putative cell-type specific lPBN subpopulations. AAV encoding a fluorescently-tagged channelrhodopsin (0.5μl, bilateral) was injected into the lPBN of the following genetic mice harboring knock-in alleles for Cre-recombinase: **(A)** Somatostatin (SST-cre, Taniguchi et al. 2011). **(B)** Calretinin (CR-cre, Taniguchi et al. 2011). **(C)** Corticotropin releasing hormone (CRH-cre, Taniguchi et al. 2011). **(D)** Neurokinin-1 receptor (NK1R-creER, Huang et al. 2016). **(E)** Neurotensin (NTS-cre, Leinninger et al. 2011). Scale bar = 200 μm (A - E).

**Figure S6.**
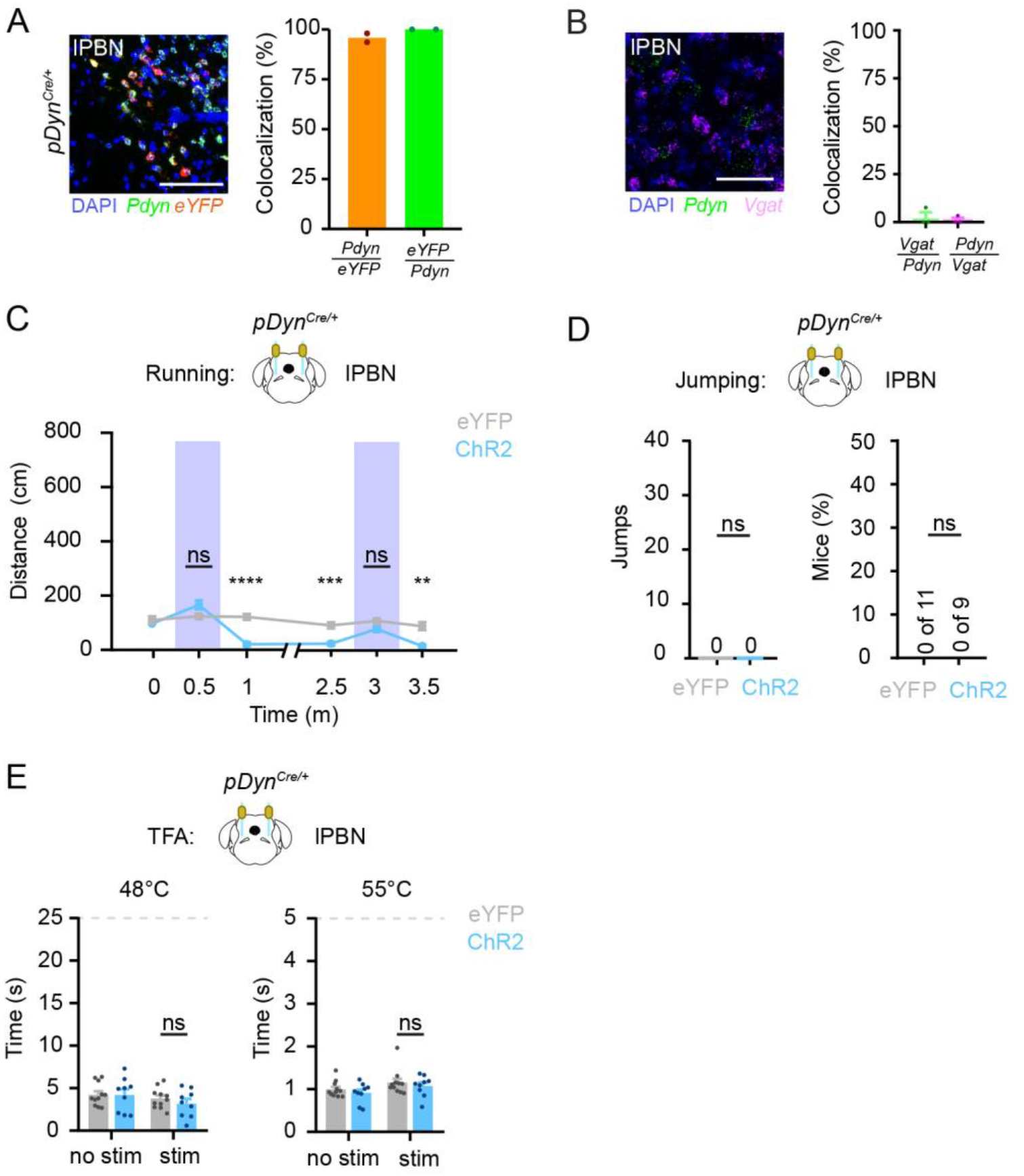
Analysis of Pdyn neurons in the lPBN and the effect of activating them. **(A)** Representative image (left) and quantification (right) of lPBN in pDyn^Cre^ mice injected with an AAV encoding Cre-dependent eYFP into the lPBN and subsequently analyzed by dual FISH using probes targeting *Pdyn* and *eYFP*. Data are mean and dots represent data points from individual animals (n = 2 mice). **(B)** Representative image (left) and quantification (right) of the lPBN using dual FISH with probes targeting *Pdyn* and *Vgat*. Data are mean and dots represent data points from individual animals (n = 3 mice). **(C)** Optogenetic stimulation of ChR2-expressing pDyn^Cre^ neurons in lPBN does not induce lateral locomotion (two-way RM ANOVA followed by Holm-Sidak post-hoc test, F (5, 90) = 17.21). **(D)** pDyn^Cre^ ChR2-injected mice do not jump when photostimulated. **(E)** Optogenetic stimulation of ChR2-expressing pDyn^Cre^ neurons in lPBN does not increase tail flick latency at 48°C (two-way RM ANOVA, F (1, 18) = 0.6300) or 55°C (two-way RM ANOVA, F (1, 18) = 0.004203) compared to control mice. Data represent mean ± SEM. Scale bar = 50 μm (A - B)

## Supplemental Methods

### Animals

The following were obtained from Jackson Laboratory: SST-cre (Taniguchi et al., 2011) stock: 013044), CR-cre (Taniguchi et al., 2011) stock: 010774), CRH-cre (Taniguchi et al., 2011) stock: 012704), and NTS-cre (Leinninger et al., 2011) stock: 017525) were obtained from Jackson Laboratories. KOR-cre was developed in the Ross lab (Snyder et al., 2018).

### Fos Induction

To induce Fos in optically implanted mice, mice were photostimulated at 10mW, 20Hz, and 5ms pulse duration for 20 minutes at a 3 seconds on, 2 seconds off stimulation pattern and subsequently perfused 90 minutes after the initial onset of photostimulation as noted for immunohistochemistry. 65um thick transverse sections of brain were collected on a vibratome and processed free-floating for immunohistochemistry as detailed in **Methods**. To quantify Fos-labeled cells, 3 optical plains separated by 10μm from the center of each section was merged into a single layer and counted for each region of interest (lPBN, BNST, CEA, VMH, and lPAG).

